# Holocephali From the Irati Formation (Paraná Basin), Brazil: Origin, Paleogographical and Paleoenvironmental Considerations

**DOI:** 10.1101/2020.10.09.333591

**Authors:** Artur Chahud

## Abstract

The Permian (Cisuralian) Irati Formation, from the Brazilian southeastern Paraná Basin bears, at some levels, Chondrichthyes, besides other vertebrates. Outcrops of this unit are frequent at the state of São Paulo eastern belt. Two members of the Irati are recognized at this state, Assistência, the upper, and Taquaral. A sandy facies, mostly at the base of the Taquaral, is noteworthy by the richness of the Chondrichthyes, mainly Holocephali. The Petalodontiformes are the Chondrichthyes most abundant, so far referred to *Itapyrodus punctatus*. Recent studies of several specimens revealed that some morphotypes must belong to different species of the genus *Itapyrodus*. Others are akin to this genus, justifying a proposition of an endemic family Itapyrodidae. The presence of this endemic family is an argument, among others, for a proposed isolation of two Brazilan Basins northeast Parnaíba and southeast Paraná, during the time of deposition of the Irati, inasmuch as Itapyrodidae are present in both basins.

## 1 Introduction

The Brazilian Paraná Basin Late Paleozoic lithostratigraphic units, present in the State of São Paulo, begin with Latest Carboniferous – Earliest Permian Itararé Group, followed by Tatuí Formation and Passa Dois Group. The earliest unit of this group is the Irati Formation (Cisuralian). This formation is distinguished by characteristic important fossil content and diversified paleoenvironments of deposition. Two members: Taquaral and Assistência (BARBOSA and GOMES, 1958; MILANI et al. 2007; HOLZ et al. 2010), make up this formation.

Two lithofacies are recognized in the Taquaral: a sandy one, mostly laid down at the base of this member and a silty one, thicker and lithologic more homogeneous. The sandy facies, even though thinner and not always present, consists of a lateral variable granulometric sandstone, fine to conglomeratic, with a great richness of isolated pieces of vertebrates, not only taxonomically different but also in number of specimens (ASSINE *et al*, 2003; CHAHUD and PETRI, 2010a).

Such a richness of fossils, was only in recent years object of detailed investigations, with their results issued in several publications (CHAHUD and PETRI 2008a, 2008b, 2014 and CHAHUD et al. 2010). The fossils already disclosed are Chondrichthyes teeth and spines, Osteichthyes teeth, scales and bone parts and tetrapods bone parts

The Holocephali were much diversified in Paleozoic, going up to the still living chimaeras (LONG, 2011). Three orders are known in the Paraná Basin; Eugeneodontiformes, Petalodontiformes and Orodontiformes (WURDIG-MACIEL, 1975; TOLEDO et al. 1997; RICHTER, 2007; CHAHUD et al. 2010).

New Holocephali specimens were recently gathered from the State of São Paulo Taquaral sandy facies, resulting in a larger and diversified fossil assemblage.

The objective of this contribution is to disclose these new forms and to propose hypothesis on their origin and paleoenvironment.

## 2 Materials and Methods

The fossils were collected from a bedding-plane exposure, along an area of approximately 7 m×20m at the base of the Taquaral Member, SW margin of the Cabeça river, about 850 m NNW from the entrance to the Santa Maria homestead, Municipality of Rio Claro, São Paulo, Brazil (UTM: 23K 0227055/7517325) (Fig. 1).

**Figure 1.**
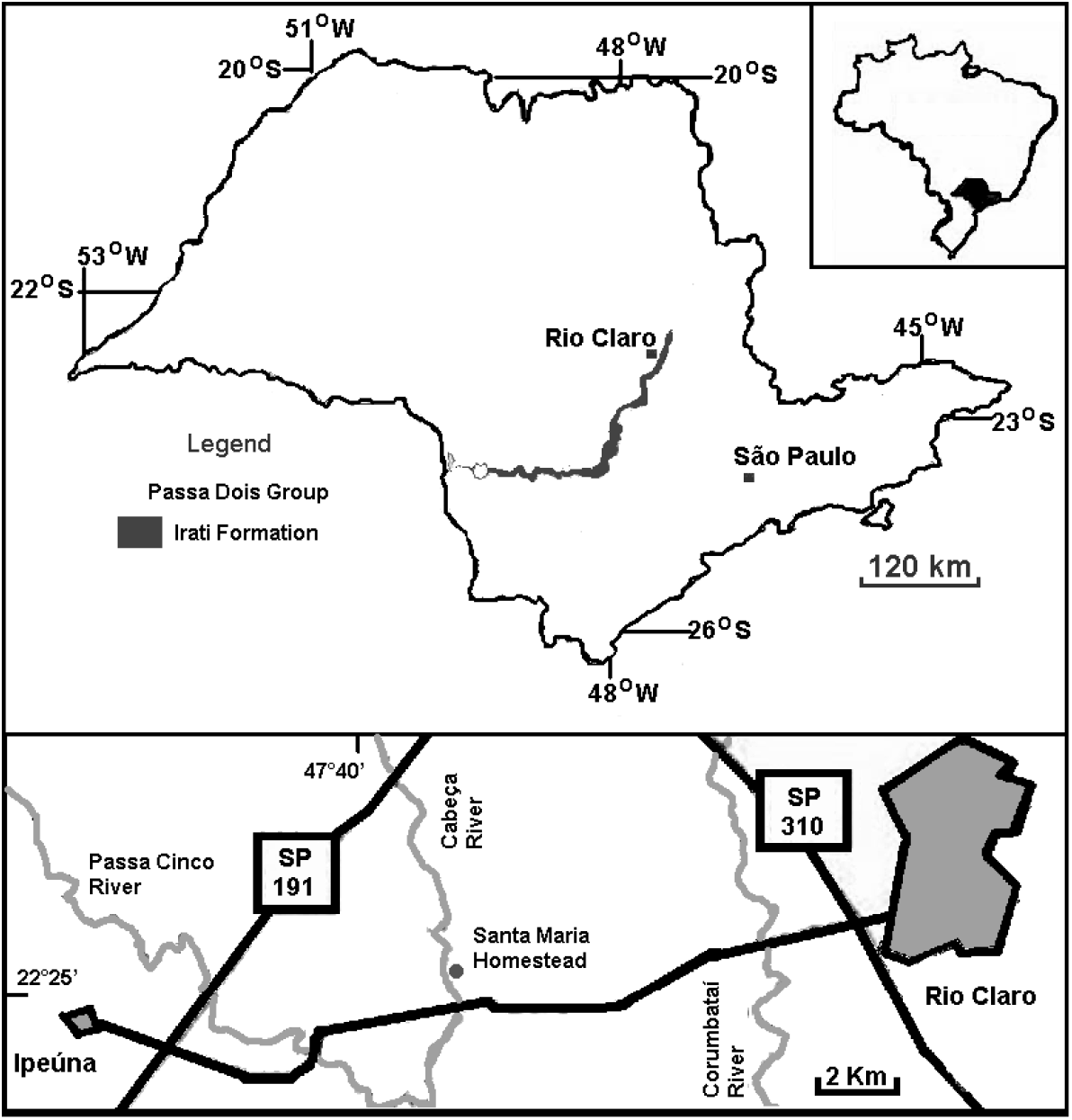
Outcrop belt of the Irati Formation in the State of São Paulo (above) and access to the fossil locality of the Santa Maria homestead (below).

They are preserved as bioclasts in a 9.5 cm-thick bed of light-gray to gray, fining-upward, cross-laminated conglomeratic sandstone, with abundant angular to rounded granules and rare pebbles of quartz and chert, dispersed in a very fine to coarse sandy matrix. Bioclasts are firmly cemented within the sandy matrix. Specimens were cleaned or removed from the rock using needles and tweezers and examined under a stereoscopic binocular microscope. The fossils are deposited in the Vertebrate Collection of the Laboratory of Systematic Paleontology (LPS) of the Institute of Geoscience of the University of São Paulo (IGc-USP).

## 3 Systematic Palaeontology

> Class Chondrichthyes Huxley 1880
>
> Subclass Holocephali Bonaparte 1832-41 (“Euchondrocephali” Lund and Grogan 1997)
>
> Order Petalodontiformes (?) Zangerl 1981
>
> Family Itapyrodidae
>
> **Genus** *Itapyrodus* **Silva Santos 1990**
>
> *Itapyrodus punctatus* **Silva Santos, 1990**

Material. GP/2E-6290; GP/2E-6304; GP/2E-6307; GP/2E-6308. Isolated teeth.

Locality. Flat-lying outcrop on the SE side of the Cabeça river (UTM: 23K 0227055/7517325), about 850 m NNW from the entrance to the Santa Maria Homestead, at the limit between the municipalities of Rio Claro and Ipeúna, São Paulo, Brazil.

Stratigraphy. Type material collected by L. I. Price in the Pedra do Fogo Formation (Permian), Parnaiba Basin, 6 km south of Pastos Bons, Maranhão, Northeast Brazil. New materials from the base of the Irati Formation (Cisuralian, Early Permian), Paraná.

Generic diagnosis (Silva Santos, 1990). Elasmobranchs, known only from teeth, showing heterodontia with distinctive symphyseal and posterior–lateral teeth. Teeth close-fitting but not fused into a dental plate. Each tooth bears a smooth crown. Symphyseal teeth bear a high crown and are labio-lingually compressed, elongate, and slightly inclined toward the lingual face; the upper margin of the crown has distinct extremities, anteriorly rounded and pointed and laterally sloping slightly downward. Posterior–lateral teeth with low, convex to nearly flat crowns. Base smooth, closed and concave. Lateral borders of the base exhibit grooves and bulges.

Specimen GP/2E-6290 (Figs. 3A-D) Labial and lingual faces of the crown disposed at a 90° angle, just below a cutting crest turned to the lingual face.

**Figure 2.**
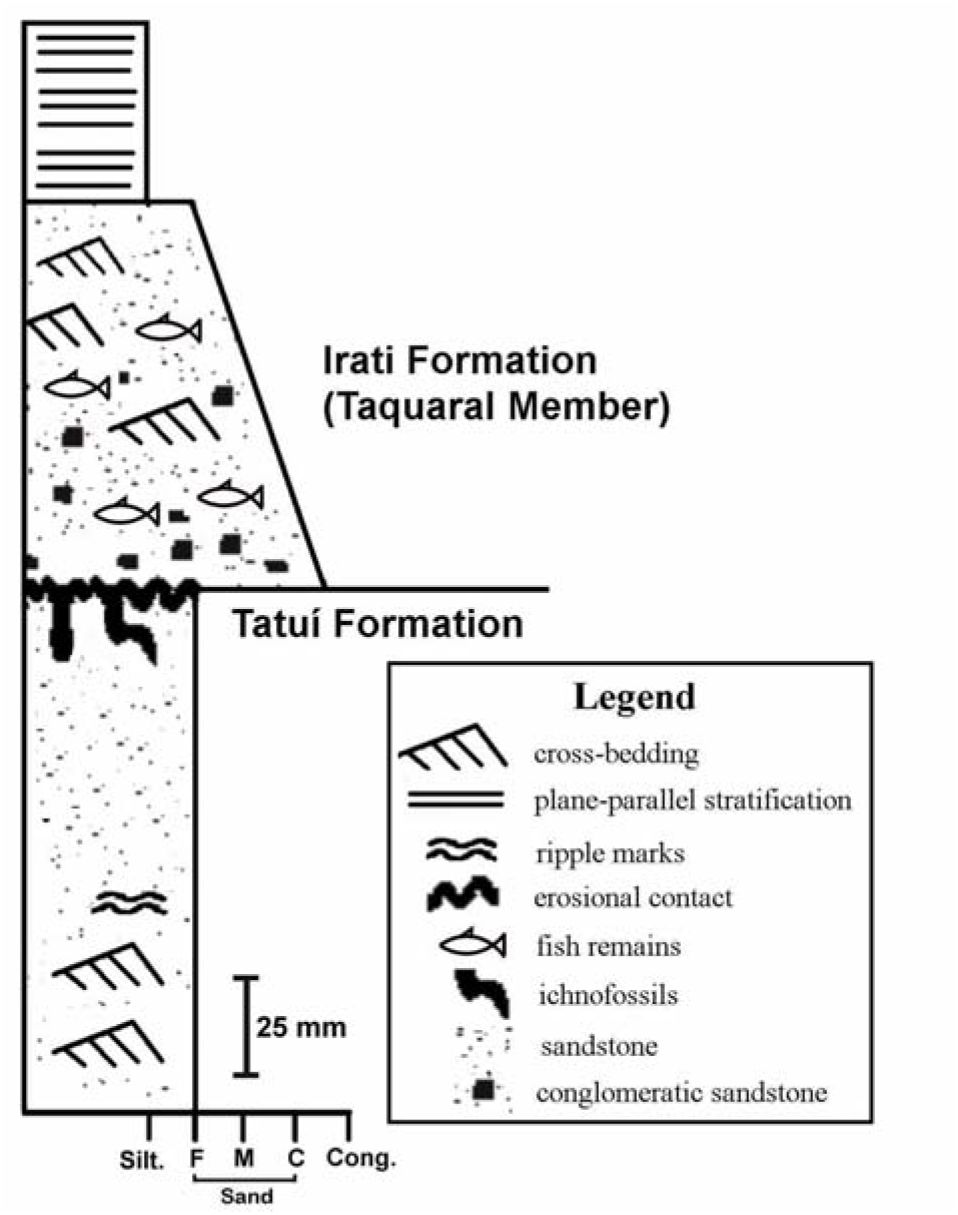
Schematic cross-section of the fossiliferous bed, homestead Santa Maria, Rio Claro, São Paulo, Brazil.

**Figure 3.**
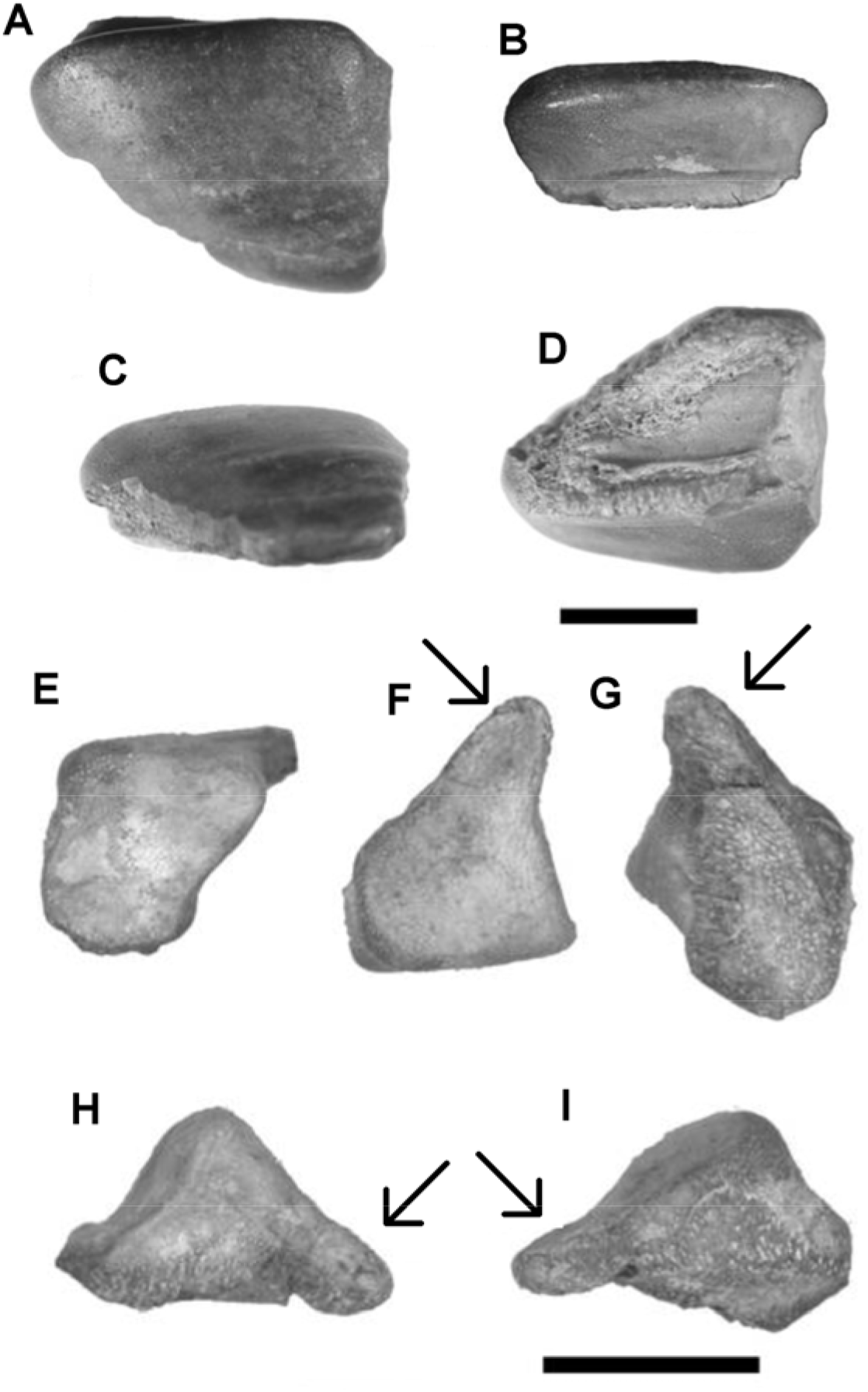
*Itapyrodus* sp. A-D) Posterior-lateral tooth (GP/2E-6290). A) labial-occlusal view; B) lingual view; C) labial view; D) inferior view. Bar scale = 5mm. E-I) Intermediate tooth (GP/2E-6258). E) lingual view; F) labial-occlusal view; G) inferior view; H) e I) laterais view. Bar scale = 10mm. Arrows at F-I; Pathological extension of the labial face.

The labial face is trapezium-shaped (Fig. 3A), but with one lateral greater. Both laterals are rather straight. The labial face is plane, with undulations on the surface (Fig. 3C) smaller in height and greater in lenght, rather worn out.

The lingual, very smaller than the labial, exhibits a small central concavity and an extension at one of the laterals (Fig. 3B).

A tip extends anteriorly, along the upper anterior lateral face, whereas a smaller extension is seen on the posterior lateral face. Small center cavities are seen on both sides.

The base articulation is plane near the lingual portion where a small cavity is seen. Crenulations are seen on the base surface (Fig. 3D).

The tooth maximum length is 13.7mm, 5mm height and 12mm width. The labial face is 8mm, labial-lingual direction and 12mm long between the laterals. The lingual is 12mm long and 5mm high.

Specimen GP/2E-6258 (Fig. 3E-I) the angle between the labial and lingual faces is 90°. The labial face is flat, extended at one of the laterals (pathology) (Figs. 3H-I)

The lingual face, 8mm high, is smaller then the labial, 11mm, exhibiting a central cavity separating the whole tooth base.

The tooth is an open curved triangle at lateral view, with one smaller side (Fig. 3H-3I).

The articulation base is irregular with very large crenulations at the labial-lingual direction, 14mm long, small at anterior-posterior direction, 5mm, measured at the lingual region and 2mm at the extended labial.

The anterior portion of both labial and lingual are rounded while the posterior portion is straight and angular. The crest is 7mm long.

The specimen is strong worn out, resulting in a clear punctuated surface on every face.

Specimen GP/2E-6304a (Fig. 4A-D) both labial and lingual faces outline an acute angle smaller than 45°, resulting in a crest turned toward a lingual face (Fig. 4D).

**Figure 4.**
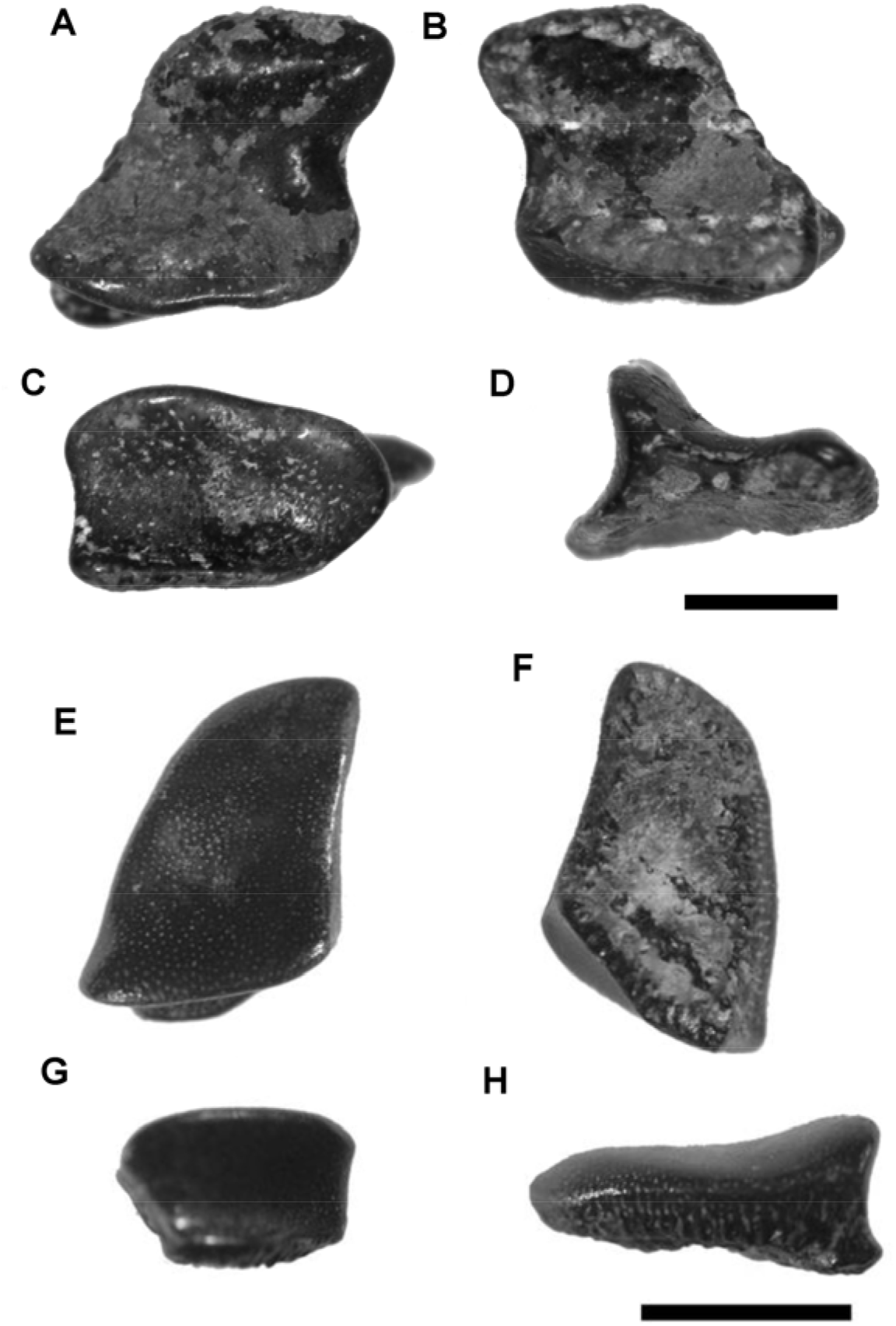
*Itapyrodus* sp. A-D) Tooth GP/2E-6304a. A) Labial – occlusal view; B) inferior view; C) lingual view; D) lateral view. Bar scale: 2mm. E-H) Tooth GP/2E-6304b. E) labial – occlusal view; F) inferior view; G) lingual view; H) lateral view. Bar scale: 2mm.

The base is straight, with irregularities and crenulations (Fig. 4B). The labial face is parallelogram shaped, with one side larger than other side, strongly curved, exhibiting a projection toward the posterior part of the tooth (Fig. 4A).

The labial face is characterized by a strong concavity near the crest and by a slight convexity at the distal extremity whereas a small concavity is present on the center part of the lingual face. The labial face is much larger, 4.5mm high, then the lingual face, 2.3mm high.

Specimen GP/2E-6304b (Fig. 4E-H) the labial and lingual faces are disposed at an angle slightly smaller than 90°, with a crest turned toward the lingual face.

The base is irregular straight and with crenulations. It is smaller then the crown width, measured at the anterior-posterior direction, with 4mm at the base and 5mm at the crown, here marked by the crest length (Fig. 4G-H).

The labial face is trapezium-shaped, with one of the sides greater, one of the tips rounded and other sharp.

The lateral borders are different at both sides, convex at the greater portion and almost straight at the smaller portion, where it exhibits a small curvature near the lingual extremity.

The labial face exhibits a strong concavity near the crest and a small convexity at the distal extremity. The lingual face exhibits a small center concavity.

The lingual face is 4mm high while the labial, 9mm. Such a striking difference puts apart this form from the others.

Specimen GP/2E-6307 (Figs. 5A-D) the lingual face is larger, 4mm long, than the labial, 2mm, measured at labial-lingual direction. The crown is strong, turned toward the lingual face, rounded at the top, with a crest disposed on the anterior-posterior lingual face. The labial face is flat and inclined toward the lingual face. This face exhibits a strong concavity, starting on the central crown, keeping stronger near the lingual border.

**Figure 5.**
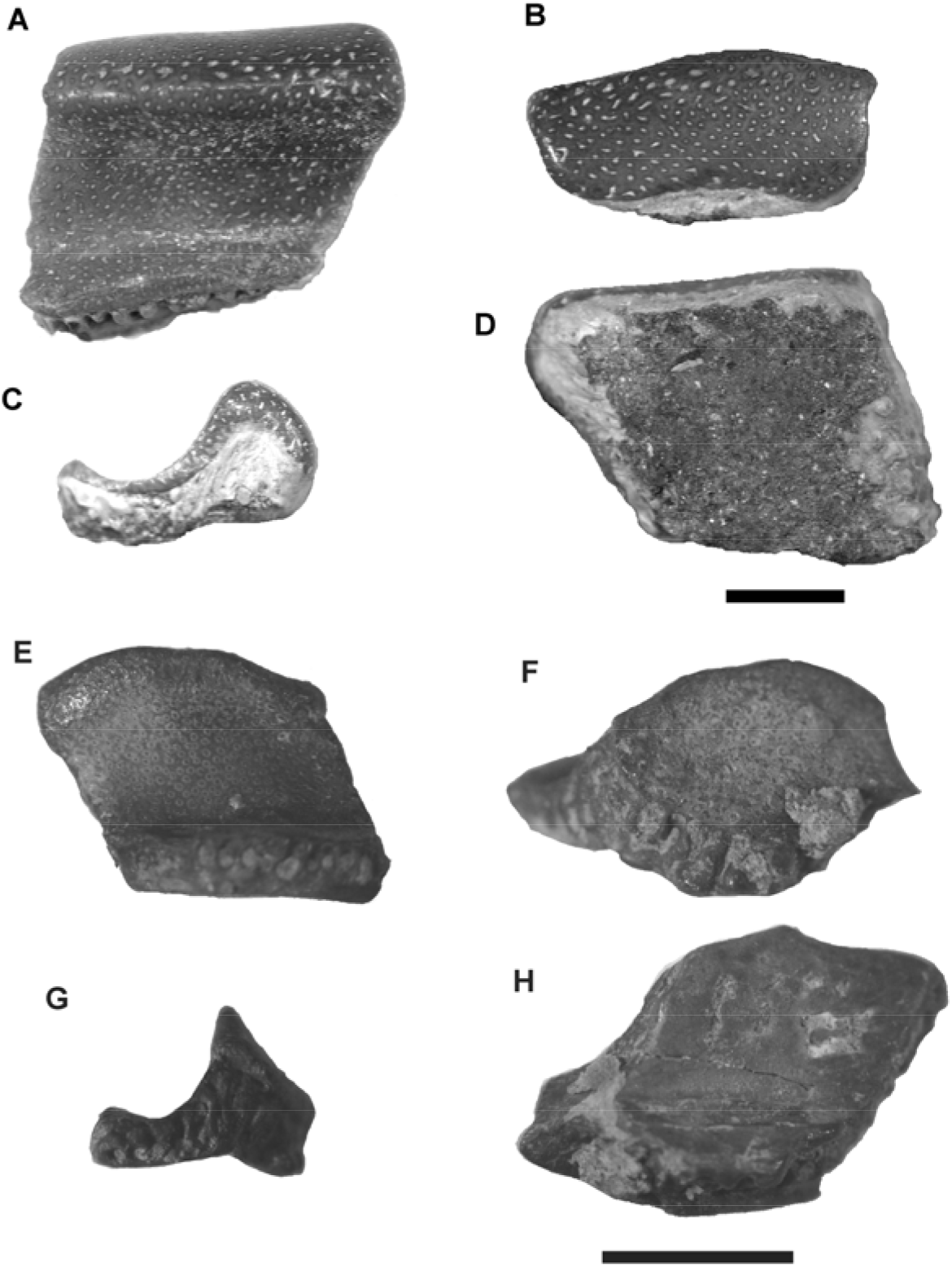
*Itapyrodus* symphyseals teeth. A-D) Tooth GP/2E-6307. A) lingual view; B) labial view; C) lateral view; D) inferior view. Tooth GP/2E-6308. E) lingual view; F) labial view; G) lateral view; H) inferior view. Bar scales 2mm.

The inferior portion of the base is flat and smooth, only with some irregularities, more clear at the lingual border. The maximum length is 7.5mm diagonally measured. The labial border is 5.5mm long and the lingual 4mm.

The crown, 4mm high, is askew toward one of the laterals, top rounded, strongly convex on its center. The crest is a little over 5mm long.

Both the anterior and posterior faces are smooth straight. The posterior one is more inclined than the labial margin.

Specimen GP/2E-6308 (Figs. 5E-H) Tooth with a depressed elongate crest, disposed along the anterior-posterior direction. The labial face is an inclined plane. The lingual face exhibits a strong concavity on the central crown.

The lingual face is 5mm wide and the labial, only 3mm, measured along the labial-lingual direction.

The smooth flat base is shaped like a parallelogram. It bears crenulations on the lingual region. The maximum length, 8mm, was measured diagonally. Both the labial and lingual borders is 4mm long.

The crown, 4mm high, is askew toward the posterior lateral. Its top is convex, increasing the convexity at the posterior portion. The crest is 5mm long.

Both lateral faces (anterior and posterior), are straight and flat, keeping the same inclination.

The specimen is worn out, allowing the clear visibility of the punctuations all over the faces.

Discussion – the first *Itapyrodus* fossil came from an outcrop in Brazilian northeastern Parnaíba Basin Permian Pedra de Fogo Formation at State of Maranhão, collected by Llewellyn Ivor Price in 1948. RAGONHA (1978) informally reported it, suggesting to be a petalodont new genus and species, *Itapyrodus punctatus*. Silva Santos (1990) described and formalized this taxon, based only on Pedra de Fogo specimens. *Itapyrodus* is a Chondrichthyes known only from teeth, showing heterodontia with distinctive symphyseal and posterior–lateral teeth.

Although none of the teeth, reported here, were found in physical contact each other, their positions within the fish mouth, may be inferred from features of their lateral faces and crests. For example, the sharp crest evidently used for nipping or cutting must be located at the front of the mouth. The lateral posterior margins teeth could easily have accommodated to the anterior bulges of adjacent tooth forming a continuous, close-fitting surface for crushing food.

The crown of the specimen GP/2E-6258 (Fig. 3E-I) is higher than a conventional postero-lateral tooth even though unlike typical symphysian. The worn out crest top, separates the lingual from the labial face, disposing diagonally between the laterals. This specimen seems to be an intermediary tooth between the symphysians and the posterior-laterals, suggesting a gradual change of dentition.

The tooth GP/2E-6290 (Fig. 3A-D) should be placed at *Itapyrodus punctatus* posterior–lateral teeth sequence, according to SILVA SANTOS (1990).

The *Itapyrodus* teeth GP/2E-6304a and GP/2E-6304b (Fig. 4), stand out as compared with other *Itapyrodus* teeth. The GP/2E-6304a is a posterior-lateral with well developed labial face, however its lingual face bears a strong salient crest. GP/2E-6304b bears an extended labial face larger than other *Itapyrodus* teeth with similar positions in the mouth. This odd anatomy may be a result of a different feeding way.

The punctuated aspect of the specimens GP/2E-6307 and GP/2E-6308 (Fig. 5) and the high crown disposition, are typical of symphysians teeth of *Itapyrodus*. However, different from *I. punctatus*, the labial face is very smaller than the lingual face. Therefore, these teeth must not be placed in the specie *I. punctatus*. The teeth are slender than those of *I. punctatus,* suggesting different feeding way, as small fishes or easy prey.

The genus *Itapyrodus*, since Silva Santos (1990) was thought to be Petalodontiformes (CHAHUD et al. 2010; RICHTER et al. 2013). As only teeth are known, the right position of this genus in this order is polemic.

According to SILVA SANTOS (1990), *Itapyrodus* was classified as Petalodontiformes, based on the following similarities with *Chomatodus, Antliodus* and *Tanaodus/Climaxodus*: the shape of the labial-lingual faces, high crowns on the symphysian teeth and, like some species of *Tanaodus/Climaxodus* (WOODWARD, 1919) and *Chomatodus* (EASTMAN, 1903) absence of teeth fusion.

SILVA SANTOS (1990) considered the taxonomic position within Petalodontidae with caution, due to the smooth base and no denticles, as is the case of *Polyrhizodus* and species of *Petalodus*, both from north hemisphere.

LUND et al. (2014a and 2014b) issued an important and pioneer systematic work, assembling genera with teeth base characteristics allowing confront with Petalodontiformes. They discard *Chomatodus* and *Tanaodus/Climaxodus* from other Petalodontiformes.

LUND et al. (2014b) suggested “Chomatodus Group”, sister group of Petalodontiformes for Chomatodus and *Tanaodus/Climaxodus* as is an important improvement for understanding the Petalodontiformes systematic. The which work, issued in 2014, of course implies that further contributions of new taxa, improve LUND et al.’s schema. So, the genus *Itapyrodus* is questionably included in the Petalodontiformes, which but discarded its inclusion in Petalodontidae, after a thorough analysis of other genera of Petalodontidae

The smooth and punctuated crown *Itapyrodus* teeth are like those of the taxa of the family Janassidae, such as *Janassa*, and Pristodontidae, similar to *Megactenopetalus* (OSSIAN, 1976; MERINO RODO and JANVIER, 1986)

The specimens dealt with herein, differ from other Holocephali like Cochliodontiformes, by the absence of teeth surface undulations, as is the case of *Deltodus, Helodus* and *Cochliodus*. They differ also by the absence of plate folds attaching on mandibular region (STAHL, 1999).

The *Itapyrodus* posterior-lateral teeth are similar to the external surface shapes of *Psammodus* and *Lagarodus* (LEBEDEV, 2008) but differ from them, by the rather higher crown and the sharp cutting borders.

Several Holocephali differences and similarities in comparison with those of northern hemisphere, justify a position against packing the Brazilian forms altogether in an unique family. The studied specimens allow grouping them in a new familiy, based both on teeth characteristics and their supposed position on the jaws.

The family Itapirodidae is endemic in the Permian Brazilian Paraná and Parnaíba basins. Examples of endemism during Taquaral deposition was disclosed before. Taquaral Member endemic taxa were recognized, by MEZZALIRA (1952) and CHAHUD and PETRI (2013a; 2013b) such as the crustaceans *Clarkecaris* (Family Clarkecaridae BROOKS, 1962) and a crustacean form still indeterminate CHAHUD and PETRI (2013b).

The Itapyrodidae appears suddenly in Paraná and Parnaíba basins in early Permian, without any register in older deposits. Originally reported in Pedra de Fogo Formation of the Parnaíba Basin, by RAGONHA (1978), in a thin but complex lithologic bed, consisting of coarse sandstone, breccia and conglomerate with silex, the same lithotypes reported by SILVA SANTOS (1990). It is noteworthy to emphasize that CHAHUD et al. (2010) reported similar lithologies containing *Itapyrodus* in the Taquaral Member sandy facies.

The deposits and the fossils are suggestive of similar paleoenvironment of both basins, probably high energy, shallow water and a strong continental influence, resulting in low degree salinity.

> *Fairchildodus rioclarensis* sp. nov.

Etymology – Genus in honour to Dr. Thomas Rich Fairchild. Paleontologist from the University of São Paulo, Brazil, and species after the town of Rio Claro, São Paulo.

Holotype - GP/2E-5929, a complete isolated tooth, but rather worn out.

Syntype - GP/2E-6459. Complete isolated tooth.

Type locality - Flat-lying outcrop on SW side of the Rio Cabeça, about 850 m NNW from the entrance to the Santa Maria homestead (UTM: 23K 0227055/7517325) at the limit between the municipalities of Rio Claro and Ipeúna, São Paulo, Brazil.

Stratigraphy - A conglomeratic sandstone layer at the base of the Taquaral Member, Irati Formation, Lower Permian, Cisuralian, immediately above the Tatuí Formation.

Material - GP/2E-5929 and GP/2E-6459. Two complete isolated teeth.

Diagnosis – Elasmobranchii known only by the teeth. High crown, without cusps, elongated sense lingual-labial. Thickness, measured at the fixed base (root) equal or smaller than the upper crown. The whole teeth lateral sides form an irregular “A”, with one lateral greater, lateral center convex in one side and concave in other side.

Row teeth touching each other but do not form dental plates. Apex longitudinal crest separates the labial and lingual faces, which form an acute angle.

Description – The GP/2E-5929 is a complete tooth, looking little worn out, “A” – shaped base bilobate and concave, with a small straight prominence in the center base, possibly related to the articulation system.

The very high crown, marked at the top by an elongate and curved longitudinal crest, sense the tooth anterior – posterior, inclined a little toward the lingual face. The crest run all along the crown, going beyond its limit to form an acute dot prominence, at the tooth posterior face.

The central concavity exposes, at the anterior face, a dentine system, from the base to the top of the crown (Fig. 6A). The tubular dentine, at the posterior face, is not so clearly seen, but still evident by the punctuated system.

**Figure 6.**
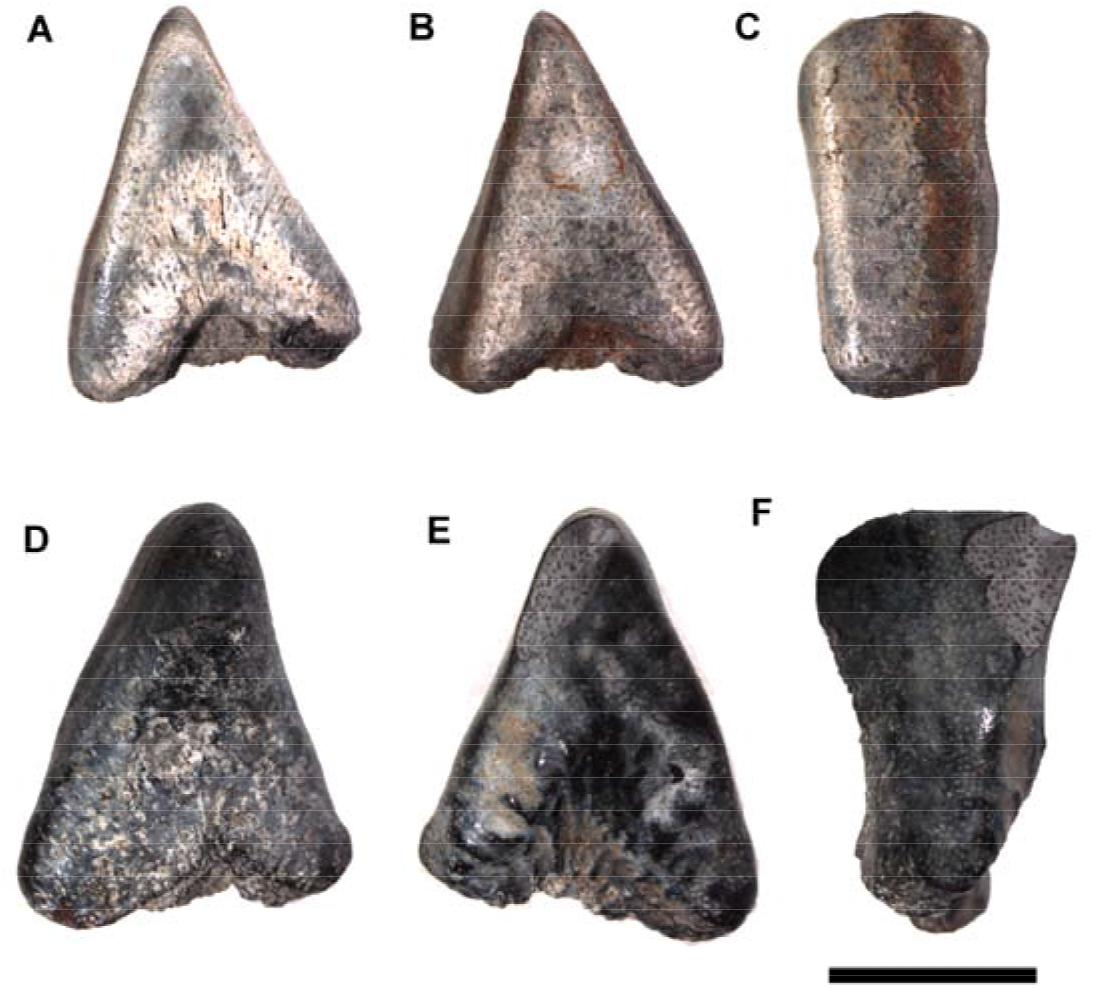
*Fairchildodus rioclarensis*. A-C) Holotype tooth GP/2E-5929: A-B) lateral views. C) lingual view. D-F) Tooth GP/2E-6459: D e E) lateral views. F) lingual view. Bar scales: 8mm.

Both lingual and labial faces are similarly flatted, resulting a rectangle (Fig. 6C), but it is still possible to distinguish the faces, by the top crest inclination, toward the lingual, the angle between the faces is about 40°. This specimen GP/2E-5929 is 14,5mm long; 10,5mm wide at the base, the crest, 7,0mm long.

The specimen GP/2E-6459, a rather larger specimen, is similar, measaring16,3mm long, 13mm wide at the base and the crest, 10,0mm long.

The two common features of the teeth are the “A” outline formed by the laterals (Figs. 6A-B and 6D-E) and differences in size of the laterals, though the difference is greater in the GP/2E-6459 (Figs. 6D-E). This specimen also is flattened on the base, exhibiting a greater central concavity at the anterior face, greater than in the specimen GP/2E-5929.

Discussion – The “A” shape also is present in species of Helodontiformes, *Helodus coniculus,* United States Mississippian e H. *appendiculatus*, British Mississippian (STAHL, 1999). However no cusps, denticle or bulge at the occlusal region and no dental plates are present in the Brazilian specimens, which discard their collocation within the Helodontiformes.

The Brazilian teeth seem to belong to similar sequences as is the case in the Eugeneodontiformes *Lestrodus, Parahelicoprion, Helicoprion, Sarcoprion, Parahelicampodus* and *Helicampodus*. However, the sequences in these genera are definied by the growth and later substitution by the spiral teeth (ZANGERL, 1981). The *Fairchildodus* specimen GP/2E-6459 might be more immature of the sequence. There are little worn out evidence by current transport. The rather sharp worn out might be caused by feeding degradation still on the jaw which would indicate absence of substitution by the tooth behind.

The Brazilian specimens, now assigned to *Fairchildodus*, were compared by CHAHUD (2007) to the Pennsylvanian *Petalodus ohioensis* from Ohio, United States, and to the Italian Alps (DALLA VECCHIA, 1988), in view of the teeth triangular shape laterals of *Fairchildodus* similar to the labial and lingual faces at the base of dental plates. However, the *Petalodus* laterals are thin or tapering to the vertex, distant from the triangle. The same occurs at the living and fossils Neoselachii with triangular teeth.

A longitudinal crest going up to the tooth apex is similar to *Itapyrodus*. However, a bulged point is present on the *Fairchildodus* teeth apex, which might mean a different feeding habitat.

The *Fairchildodus* teeth central lateral faces are concave at one side and convex at the opposed correspondent side. This situation suggests a position in sequence on the jaw, forming two lateral rows, as SILVA SANTOS (1990) suggested for *Itapyrodus*. There would be no internal rows or palatal teeth.

The *Fairchildodus* teeth are not robust like those of the symphysean teeth *Itapyrodus* (Fig. 7) so they are less fit to break food but fit to cut. It is inferred therefore a different mandible and so, a different feeding way.

**Figura 7.**
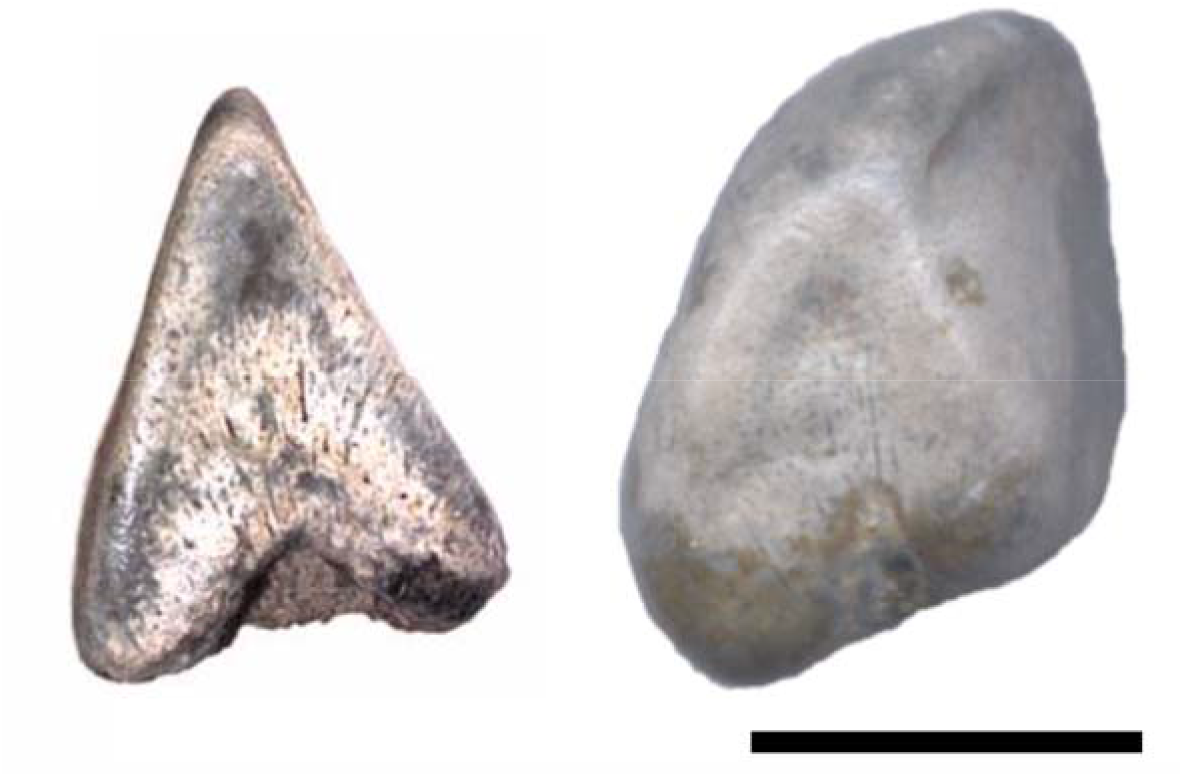
Comparison between the sides of a *Fairchildodus* and *Itapyrodus*, especially the form of "A". Bar scale: 10mm

The A-shaped lateral faces are similar in both genera (Fig. 7). However the labial and lingual faces are very evident in *Itapyrodus* and discreet in *Fairchildodus*, justifying the proposition of a new genus, but both in the family Itapyrodidae.

*Itapyrodus* is rather common in the sandy Taquaral facies while *Fairchildodus* is rare. It is believed then that this last genus was derivate from *Itapyrodus* inside the Paraná Basin and exclusive of this basin.

## 4 Taphonomical and Paleoenvironmetal Considerations

The conglomeratic sandstones depositional archictecture of the Taquaral Member sandy facies means a high energy water deposition. CHAHUD (2011) called attention for the bad-sorted deposits, the fossils reinforce this interpretation. Most of them are fragmented, but found together with articulated good preserved fossils in the same beds. As a whole, the fossil dispositions are chaotic as seen from above.

Finning upward from a coarse bad sorted sandstone with more than 1cm clasts to fine sandstone, mean a transgression with gradual increase of water depth.

The fossils were submitted to little exposition before setting on the ground, as inferred by the following observations: A) some teeth are in good state of preservation with no or very small flaws (CHAHUD and PETRI, 2009, 2010a, 2010c). B) great fragmented “amphibious”, still keeping some kind of articulations and teeth still connected with jaw parts (CHAHUD and PETRI, 2010b). It is inferred therefore, small reworking or small exposition of the fossils before setting down on the ground. Older reworking deposits or long time of wave action (ASSINE et al. 2003) are, then, discarded.

The different state of preservation of the fossils, might be due to water energy sudden changes, caused by higher energy short episodes, taking the deposits to points of better chance of preservations.

A coast with strong continental influence is the best fit for one that generated this deposit. Besides, MARASCO et al., (1993) reported few acritarch correlated beds with the ones here studied. These acritarchs would eventually indicate some salinity but not so great to allow the presence of exclusively marine organisms.

The occurrence of characteristic fluvial fossils (basal tetrapods), the scarce presence of acritarchs and the absence of stenohaline species, discard ocean normal salinity paleoenvironment.

The most convinced hypothesis for the paleoenvironment of deposition of the lowermost sandy Taquaral beds, was a large water body, under strong continental influence and low salinity, as are the cases of the Baltic Sea, Black Sea and Caspian Sea, this completely isolated. The Baltic Sea, when completely isolated, resulted in lacustrine with low salinity deposits and few acritarch taxa (BRENNER, 2005).

## 5 Origin and Evolution of Paraná Basin Holocephali

Chondrichthyans are known in the Paraná Basin, since the Pennsylvanian (BARCELLOS, 1975) and Early Permian (SILVA SANTOS, 1947) older than Irati Formation. However there is no Holocephali evidence from the Paraná Basin deposits older than Taquaral Member.

It was discarded then, the hypothesis of its origin from older taxa within this basin. On the other hand, the presence of the petalodont *Itapyrodus punctatus* and the Xenacanthiformes *Taquaralodus albuquerquei* both in the sandy facies of the Taquaral and coeval Pedra de Fogo Formation in the Brazilian northeastern Parnaíba Basin (SILVA SANTOS, 1946; RAGONHA, 1978; CHAHUD and PETRI, 2008a; CHAHUD and PETRI, 2010b, e CHAHUD et al. 2010), both endemic in these two basins and not found elsewhere, neither in Brazil nor abroad, favored the proposition of a little episode ligation of these basins in the Cisuralian, as suggested by CHAHUD and PETRI (2014). This hypothesis is reinforced by the Transbrasiliano, a Precambrian lineament northeast – southwest, since the Brazilian northeast to Argentinian Patagonian (CORDANI et al. 2010). This Precambrian structure was reactivated several times within the Phanerozoic. There is evidence of possible Silurian and Pennsylvanian (glacial) in an Agua Bonita “Basin”, midway between Paraná and Parnaíba basins, along this lineament (AGUIAR et al. 2013; CHAMANI et al. 2013). However there are no register of coeval Irati or Pedra de Fogo beds in this “basin”. These deposits, might have been eroded after deposition.

The ichthyofossils from Santa Maria Homestead sandy Taquaral facies outcrops are Permian, in spite of some taxa reminding those of the Pennsylvanian Northern Hemisphere, including among them, the genera *Orodus*, *Sphenacanthus* and *Amelacanthus*, though these taxa are also present in Permian beds (CHAHUD et al. 2010; CHAHUD and PETRI, 2014).

The sandy Taquaral beds Holocephali are the oldest in the Gondwana, which may be coeval with the Holocephali from Bolivia Copacabana Formation (MERINO RODO and JANVIER, 1986) and Australia Wandagee Formation (TEICHERT, 1940). The beds from these two formations were laid down in tropical marine paleoenvironments (HILL, 1942; MERINO RODO and JANVIER, 1986; SAKAGAMI, 1995), suggestive that these Chondrichthyes thrived only in tropical water paleoenvironments. If this assumption is right, the absence of Holocephali in the Brazilian Late Paleozoic beds, older than Taquaral, might be due to glacial and temperate paleoenvironments, unfit climates, for those fossils.

Chondrichthyes fossils are also present in the Permian younger than Irati, Teresina Formation (Würdig-MACIEL, 1975; RICHTER, 2004a; 2004b; 2007) and Corumbataí Formation (TOLEDO et al. 1997). Those from Teresina are Eugeneodontiformes and Orodontiformes, however those from Corumbataí are thought to be Petalodontiformes, similar to *Itapyrodus*. They must be offsprings from Irati times confined to the northern Paraná Basin Corumbataí.

The Corumbataí Petalodontiformes are poorly known, poorly identified. If this identification is correct, they are among the youngest register of this group.

## 6 Conclusions

Detailed researches on the Irati Holocephali revealed a diversified forms of *Itapyrodus*, many still demanding more investigations. Anyway, this genus is not monospecific.

It differs from other genera and families, known in the northern hemisphere, as a consequence of the Paraná and Parnaíba basins isolation in those times. The time of confinement was sufficient to developed an endemic family, Itapyrodidae.

A new genus and specie based on teeth, endemic in the Paraná basin, is the herein proposed *Fairchildodus rioclarensis*. Some morphologic characteristics are similar to those of *Itapyrodus,* allowing its allocation in the family Itapyrodidae. The genus *Fairchildodus* is located in Itapyrodidae based, among others, on the differences between the labial and lingual lateral faces.

No ocean fossils were recognized in the Sandy Taquaral facies. The continental influence, as fluvial floodings and storms increasing the current overflows are suggestive that Chondrichthyes thrived at low salinity and high energy paleoenviroments.

The northern hemisphere Paleozoic Petalodontiformes thrived in warm or tropical waters, both in the Carboniferou and in the Permian. On the other hand, during the Gondwana Carboniferous and earliest Permian times, the paleoclimates were cold or glacial, not appropriated for Holocephali. The Petalodontiformes, as *Itapyrodus*, must be considered good warm paleoclimate indicators, unfit to extreme cold winters.

## Acknowledgements

The authors thank the National Council for Scientific and Technological Development (CNPq) (Grant: 500755/2013-2) for financial support and the owners of the Santa Maria homestead, Luiz and Bernadete Esposti. The authors thank also the ‘‘Departamento de Geologia Sedimentar e Ambiental of the Institute of Geoscience of the University of São Paulo’’ which allowed the preparation of fossils in its laboratories.

## Notes

### Competing Interest Statement

The authors have declared no competing interest.

